# Revision of commonly accepted Warburg mechanism of cancer development. Redox-sensitive mitochondrial cytochromes in breast and brain cancers by Raman imaging

**DOI:** 10.1101/2021.02.03.429508

**Authors:** Halina Abramczyk, Jakub Maciej Surmacki, Beata Brozek-Pluska, Monika Kopeć

**Affiliations:** Lodz University of Technology, Institute of Applied Radiation Chemistry, Laboratory of Laser Molecular Spectroscopy, Wroblewskiego 15, 93-590 Lodz, Poland

**Keywords:** optical biopsy, Raman spectroscopy and imaging, brain and breast cancer, cytochrome c, cell cultures

## Abstract

**Background:** We studied oncogenic processes that characterize human breast cancer (infiltrating ductal carcinoma (IDC)) and human brain tumors: glioma, astrocytoma and medulloblastoma based on the quantification of cytochrome redox status by exploiting the resonance-enhancement effect of Raman scattering.

**Methods:** We used Raman imaging to monitor changes in the redox state of the mitochondrial cytochromes in ex vivo human brain and breast tissues surgically resected specimens of human tissues and in vitro human brain cells of normal astrocytes (NHA), astrocytoma (CRL-1718), glioblastoma (U87-MG) and medulloblastoma (Daoy), and human breast cells of normal cells (MCF 10A), slightly malignant cells (MCF7) and highly aggressive cells (MDA-MB-231) by means of Raman microspectroscopy at 532 nm.

**Results:** We visualized localization of cytochromes by Raman imaging in the major organelles in cancer cells. We demonstrated that the “redox state Raman marker” of the ferric low spin heme in cytochrome c at 1584 cm^−1^ can serve as a sensitive indicator of cancer aggressiveness. We compared concentration of reduced cytochrome c and the grade of cancer aggressiveness in cancer tissues and single cells and specific organelles in cells: nucleous, mitochondrium, lipid droplets, cytoplasm, and membrane.

**Conclusions:** We found that the concentration of reduced cytochrome c becomes abnormally high in human brain tumors and breast cancers in human tissues. Our results suggest that the mechanisms controlling the electron transport chain are spectacularly deregulated in cancers and indicate that electron transport, organized in terms of **electronegativity**, is inhibited between complex III and cytochrome c for isolated cells in vitro and between cytochrome c and complex IV in brain and breast tissues. The results provide evidence that the extracellular matrix and interactions with cell microenvironment play an important role in the mechanisms controlling the electron transport chain by cytochrome c. Our results reveal the universality of Raman vibrational characteristics of mitochondrial cytochromes in metabolic regulation in cancers that arise from epithelial breast cells and brain glial cells.

## Introduction

It is becoming clear that oncogenic processes during cancer development are governed by balance between the need of the cell for energy supply with its equally important need for macromolecular building blocks and maintenance of redox balance.^1–4^

Regarding macromolecular building blocks the role of fatty acids as critical bio-energetic substrates within the glioma cells^5–12^ and breast cells^9,13,14^ has been recognized. The redox balance that depends to a large extend on mitochondrial functionality in electron transfer chain have been extensively studied.^15–18^

Cytochrome family, a heme-containing protein, play a critical role in mitochondrial mechanism of cell respiration as an electron carrier in the electron transfer chain in mechanism of oxidative phosphorylation. Cytochromes are also important in intercellular cell signalling, metabolism of polyunsaturated fatty acids, and apoptosis. Cytochrome c is released into the extracellular space and can be easily measured in the serum serving as a marker of severe cellular damage and death. Elevated levels of serum cytochrome c has been reported in many diseases, including cancer.^18^ The results on the role of cytochrome c in brain disorders of different research groups are somewhat conflicted regarding the respiratory chain in glioma.^15,16^ Early studies on glioma cell rat xenografts identified lower enzyme expression of Complex IV, also known as cytochrome c oxidase (COX) and Complex II, (SDH) enzyme expression in more hypoxic areas of the tumor. More recently, one group reported significantly lower activity of complex II-IV in anaplastic astrocytomas and lower activity of complex I-IV in glioblastomas compared with normal brain tissue.^15^ Another group studied human glioma tissue samples by mass spectrometry and observed lower expression of some subunits in complex I, but higher levels of many oxidative enzymes including catalase.^16^

The significance of mitochondrial dysfunctionality has not been studied by Raman methods, to the best of our knowledge, but traditional biological techniques were used to study invasive ductal carcinoma (IDC). ^19^

However, further studies are required for supporting this role for cytochrome c and the responsible pattern recognition receptor(s) remain to be discovered.

Cytochromes are classified on the basis of their lowest electronic energy absorption band in their reduced state. Therefore, we can distinguish cytochrome P450 (450 nm), cytochrome c (550 nm), cytochromes b (≈565 nm), cytochromes a (605 nm). The cytochromes are localized in the electron transport chain in the complex known as complex III or Coenzyme Q – Cyt C reductase, sometimes called also the cytochrome bc_1_ complex (cytochrome b, cytochrome c1). Cytochrome c, which is reduced to cytochrome c ^Fe+2^ by the electron from the complex III to complex IV, where it passes an electron to the copper binuclear center, being oxidized back to cytochrome c (cyt c Fe^+3^). Complex IV is the final enzyme of the electron transport system. The complex IV contains two cytochromes a and a3 and two copper centers. Using different excitations being in resonance with the absorption bands of the cytochromes one can spectrally isolate various cytochrome chromophores in the complex III, cytochrome c and complex IV by Resonance Raman enhancement scattering. Using different excitations, we are able to monitor contribution of cytochrome family in the electron transfer chain in the mitochondrial respiration originating from cytochrome b, c1 in complex III, cytochrome c, and a, a3 cytochromes in complex IV. The spectral features of cytochromes a and a3 in complex IV can be observed at resonance conditions of 420 nm and 445 nm for the oxidized and reduced state, respectively as well as at pre-resonance conditions. The spectral features of cytochrome c can be monitored at 532 nm due to absorbance band (Q band) centered at 530 nm. ^20^

Monitoring redox state of mitochondrial cytochromes has been demonstrated as a versatile clinical diagnostic tool with numerous successful reports on the detection of cancerous tissues in human patients.^19,21–24^

It is evident that the real progress in improved cancer therapy and treatment depends on much better than today understanding biological mechanisms of mitochondrial dysfunction in cancer. To achieve this goal there is an urgent need to improve the conventional methods of molecular biology (immunohistochemistry, real-time PCR, immunoblotting, measurement of mitochondrial membrane potential, cell proliferation assay and caspase 3 activity assay under hypoxic conditions^19^) in cancer diagnostics and develop new imaging tools, because current imaging methods are often limited by inadequate sensitivity, specificity, spatial and spectral resolutions.^25^

Our goal was to demonstrate possibility of monitoring redox state changes occurring in mitochondrial cytochromes in cancers by Raman imaging and correlate the concentration of cytochrome c with cancer aggressiveness. It is well known that cytochrome c is undoubtedly one of the most prominent molecules in the electron transport chain required to fuel life via ATP, but its role in molecular mechanisms associated with an aggressive phenotype of cancer remain largely unclear. Better comprehension of biochemical pathways in two different types of cancer - human brain tumors and ductal cancer cells can be obtained by integrated approach consisting of in vitro cells, ex vivo tissues, and relevant biological models in vivo.

Therefore, this paper presents a truly unique landscape of cancer modern biochemistry by a non-invasive Raman approach to study redox status of cytochromes in brain and breast cancer in ex vivo surgically resected specimens of human brain and breast, normal and cancerous tissues and in vitro human brain and breast cells.

We study human normal and cancer tissues, in vitro human brain cells of normal astrocytes (NHA), astrocytoma (CRL-1718), glioblastoma (U87-MG), medulloblastoma (Daoy), and human breast normal cells (MCF 10A), slightly malignant cells (MCF7) and highly aggressive cells (MDA-MB-231) by means of Raman microspectroscopy, at 532 nm.

In this paper we explore a hypothesis involving the role of reduction-oxidation pathways related to cytochrome c in cancer development.

Understanding of cytochrome role in brain and breast cancers with new methods will help establish Raman spectroscopy as a competitive clinical diagnosis tool for cancer diseases involving mitochondrial dysfunction and is a prerequisite for successful pharmacotherapy.

## Results

### 1. Cytochromes in cancer human tissues

To properly address redox state changes of mitochondrial cytochromes in brain and breast cancers by Raman spectroscopy and imaging, we systematically investigated how the Raman method responds to in vitro human cells and ex vivo human tissues.^26,27^ In vitro experiments will allow to study a single cell by eliminating the cell-to cell interactions. The ex vivo human tissue experiments will extend our knowledge on the influence of environment on cancer development.

Recently we compared the Raman spectra of the human breast and brain cancer tissues using different laser excitation wavelengths.^27-29^ This approach might generate Raman resonance enhancement for some tissue components that cannot be visible for non-resonance conditions and provide selective spectral isolation of components crucial for understanding mechanisms of mitochondrial dysfunctions associated with cancers.

Fig. 1 shows the spectrum of isolated cytochrome c compared with the average Raman spectrum of *ex vivo* surgically resected specimens human brain tissue at the G4 level and human breast tissue at the level G3 of malignancy at 532 nm. The average Raman spectrum was obtained for 13 brain patients and 5 breast patients. For each patient diagnosed with brain cancer 6500 Raman spectra were recorded, for patients diagnosed with breast cancer the number of single Raman spectra was 8000.

**Figure 1.**
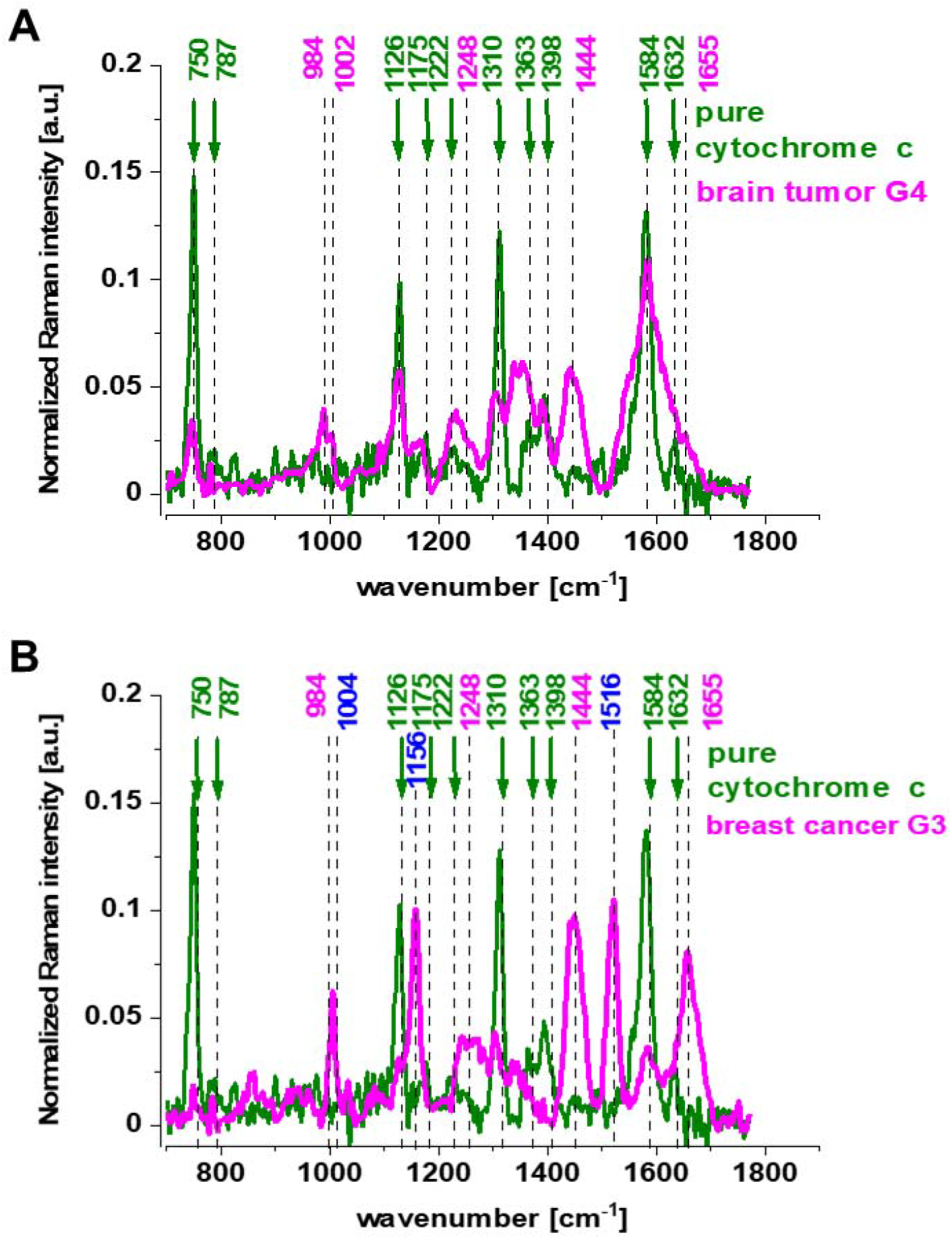
The average (number of patients n (G4)=13, number of single Raman spectra 85000) Raman spectra of the *ex vivo* human tissue of brain tumor, medulloblastoma (G4) ─,cytochrome c ─ at the excitation at 532 nm (A), the average (number of patients n(G3)=5, number of single Raman spectra n= 40 000) Raman spectrum of the *ex vivo* human tissue of breast cancer, infiltrating ductal carcinoma (IDC)) (G3) ─, cytochrome c ─ at the excitation at 532 nm (B).

The excitation at 532 nm enhances two types of components of the tissue: carotenoids (1520 cm^-1^ and 1157 cm^−1^) and cytochromes c and b (750, 1126, 1310, 1337, 1363, 1584, 1248, 1352 and 1632 cm^−1^).^30,31^ As the enhancement of carotenoids was discussed in our laboratory in many previous papers^13,32-36^, here we will concentrate on cytochrome family. Using 532 nm laser excitation one can monitor spectral features of complex III and cytochrome c due to Q bands at 500-550 nm related to intra-porphyrin transitions of the heme group in cytochrome c.^37,38^

Comparison of the Raman resonance spectra of the brain and breast tissues in Fig.1 with isolated cytochrome c shows that most vibrations in the tissue correspond perfectly to vibrations of cytochrome c^30,31^ upon 532 nm excitation. Correlation analysis shows perfect agreement between the tissue and cytochrome c peaks (Pearson correlation coefficient is 0.99951 at the confidence level 0.95 with the p-value of 0.0001).

One can see from Fig.1 that the vibrational mode at 1584 cm^−1^ appearing in the Q-resonant Raman spectrum at 532 nm is the most prominent band. The band at 1584 cm^−1^ appearing in the Q-resonant Raman spectra is characteristic of cytochrome c.^31^ Since Raman scattering is much weaker in the oxidized form of cytochrome c the peak at 1584 cm^−1^ originates from the reduced form of cytochromes c.^39^

Our results in Fig. 1 demonstrate that the Raman intensity of 1584 cm^−1^ peak is the most sensitive vibration of redox status in the cell and is related to concentration of the reduced cytochromes c. Therefore, we used 1584 cm^−1^ to study the redox state of mitochondrial cytochrome c in brain and breast cells in vitro, and in ex vivo human tissues. The Raman spectra in Figure 1 do not provide information about distribution of cytochrome c in the cancer tissues. To learn about the distribution of cytochrome c we used Raman imaging.

Fig. 2 shows the Raman spectra and images for the human brain tissue of medulloblastoma (G4) and human breast tissue (infiltrating ductal carcinoma (IDC), (G3). Creating Raman images by using specific filters for proteins, lipids, and cytochrome c as presented in Fig. 2 one can analyze distribution of proteins (red color), lipids profiles (blue color), and cytochrome (green color) and learn about the biochemical composition from the corresponding Raman spectra.

**Figure 2.**
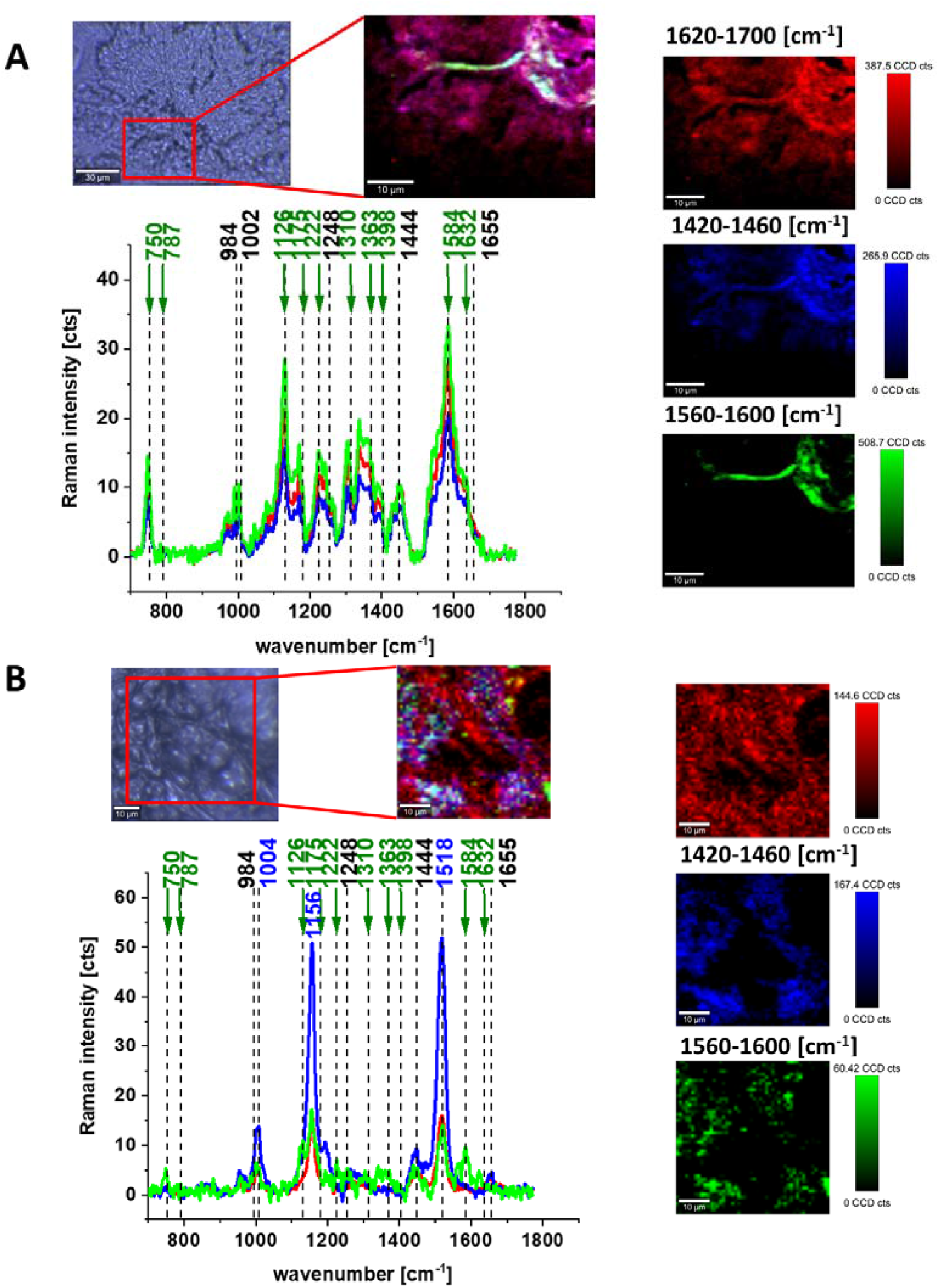
Microscopy image, Raman images of distribution of proteins (red color), lipids and carotenoids (blue color) and cytochrome c (green) for brain tumor, grade IV medulloblastoma (40×40 µm, resolution 0.5 µm, integration time 1.0 sec, the average Raman spectra of lipids – blue, proteins – red, cytochrome – green at 532 nm (A), Microscopy image, Raman images of distribution of proteins (red color), lipids (blue color) and cytochrome c (green) for breast cancer, grade I ductal cancer (40×40 µm, resolution 0.5 µm, integration time 1.0 sec, the average Raman spectra of lipids and carotenoids – blue, proteins – red, cytochrome – green at 532 nm (B).

Cytochrome Raman peaks observed for the spectra recorded at 532 nm excitation show resonance enhancement due to the electronic Q band absorption for both c and b-type of cytochromes. It has been reported ^4,5,23,43^ that the peaks at 750 and 1126 cm^−1^ are typical for both cytochrome c and b, while the peaks at 1310, 1398 correspond to cytochrome c, and 1300, 1337 cm^−1^ have been assigned to b-type of cytochromes. It means that the peak at 1337 cm^−1^ can be used to track the reduced cytochrome b form (ferrous (Fe^2+^) cytochrome). Therefore, the peaks at 750 and 1126 cm^−1^ observed in Raman spectra of human brain and breast tissues presented in Figure 3 will be used to evaluate contributions of cytochrome b and cytochrome c. The ratio of 750/1126 was used to estimate the relative contribution between reduced cytochrome c, c1 vs. reduced cytochrome b.

**Figure 3.**
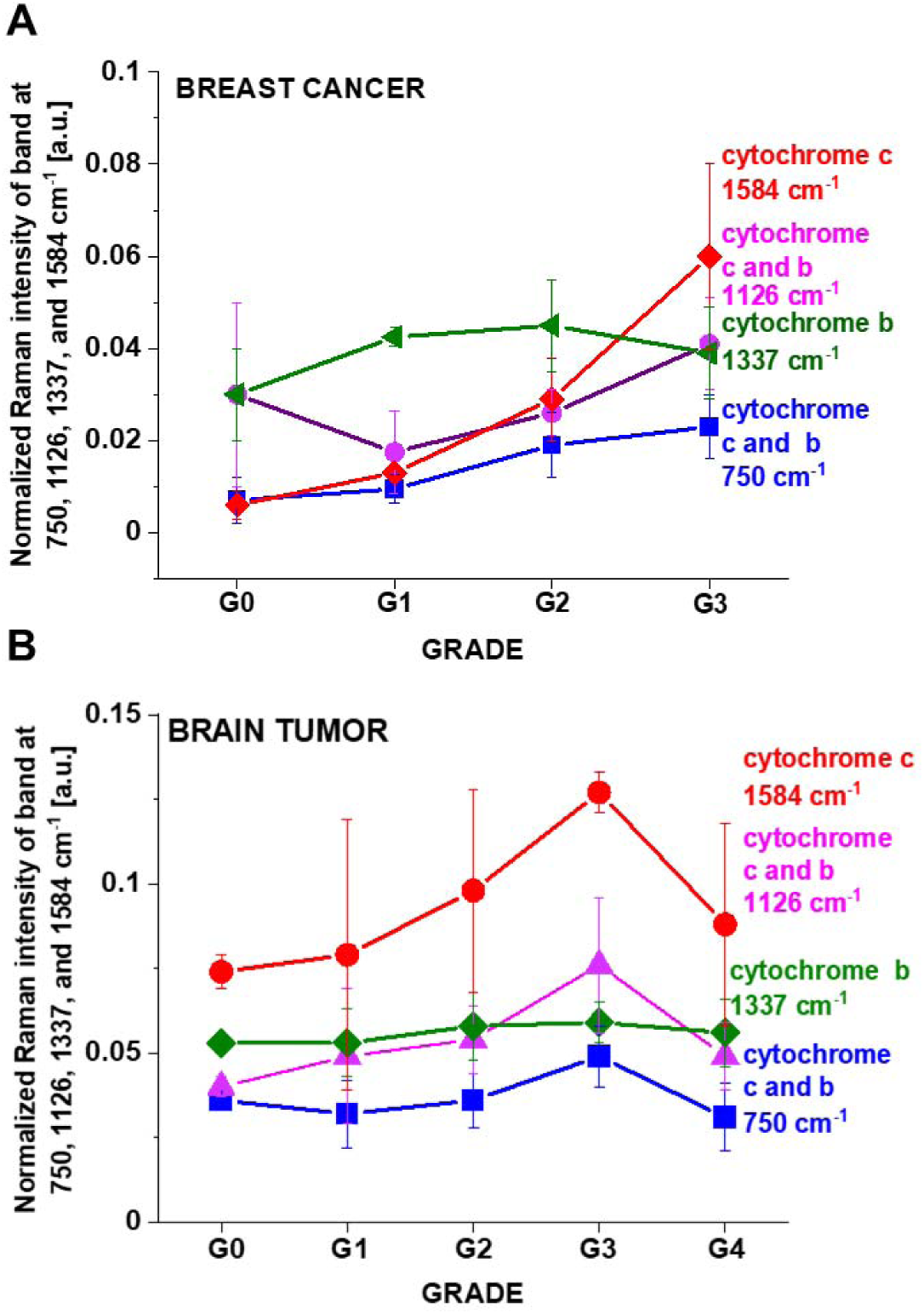
Raman intensity of the 750, 1126, 1337 and 1584 cm^−1^ as a function of grade for breast normal (G0) and cancer (invasive ductal cancer) human tissue (G1, G2, G3) (average+/-SD from number of patients n=39, number of single Raman spectra =305 000) (A), Raman intensity of the 750, 1126, 1337 and 1584 cm^−1^ as a function of grade for brain normal (G0) and tumor tissues (G1, G2, G3, G4) (average+/-SD from number of patients n=44, number of single Raman spectra = 280 000) (B).

To track the amount of the ferrous cytochrome c in brain and breast cancer tissues we have analyzed the intensity of the band at 1584 cm^−1^ assigned to the vibrational mode *v*_19_. In this spectral region there are several overlapping Raman bands (*v*_19_ of ferric heme c (1582 cm^−1^), *v*_19_ of ferrous heme c (1582 cm^−1^), *v*_2_ of ferric heme c (1585 cm^−1^), *v*_19_ of ferrous heme cyt b (1586 cm^−1^) and *v*_2_ of ferrous heme b (1583 cm^−1^). Fortunately, all vibrational modes of ferric cytochrome can be eliminated from the further discussion due to the fact that the intensities of these peaks are very weak compared to the signals of ferrous forms with one exception of the band 1634 cm^−1^. Thus, the bands at 1584 cm^−1^ (reduced cyt c) and at 1634 cm^−1^ (oxidized cyt c) of cytochrome c can be used as a very important parameter controlling the level of reduction in cancer tissues and cells.

In the view of the results presented so far we can state that Raman spectroscopy can be used in probing biochemical concentrations of cytochromes in normal and cancer cells.

To check whether cytochromes are upregulated in brain and breast cancers we studied the Raman signals corresponding to cytochromes as a function of cancer malignancy using grades. The grade is evaluated on the basis how much the cancer cells look like normal cells and measure cell anaplasia (reversion of differentiation). The malignancy for breast cancer is expressed on a three point scale G1-G3, while for brain human tumors on a four point scale G1-G4.^43^

Fig. 3A shows the intensity of the 750, 1126, 1337 and 1584 cm^−1^ Raman vibrations as a function of grade for breast normal (G0) and cancer (invasive ductal cancer) human tissue (G1, G2, G3). We calculated the average Raman spectra for each Raman vibration. The total number of patients n is 39, for each patient thousands spectra obtained from Raman imaging were used for averaging. Based on the average values obtained for the Raman biomarkers of cytochrome c and b we obtained a plot as a function of breast grade malignancy.

One can see from Fig. 3 A that the intensity of Raman biomarker at 1584 cm^−1^ corresponding to concentration of cytochrome c increases with breast cancer aggressiveness. The intensity of Raman biomarker at 1337 cm^−1^ corresponding to concentration of cytochrome b does not change with breast cancer aggressiveness. The intensities of Raman biomarkers at 750 cm^−1^ and 1126 cm^−1^ corresponding to concentration of cytochromes c and b increase with breast cancer aggressiveness.

Fig. 3B shows that for brain tissues the Raman intensities of the characteristic vibrations of cytochrome c (1584 cm^−1^) and cytochromes c and b (750 cm^−1^ and 1126 cm^−1^) first increase with increasing of cancer aggressiveness up to G3 and then decreases for G4.

The Raman intensity of the band 1337 cm^−1^ corresponding to the concentration of the reduced cytochrome b does not change vs tumor brain aggressiveness when compared with the normal tissue.

In the view of the results presented in Fig.3 it is evident that the Raman biomarker I_1584_ measuring contribution of cytochrome c in the human tissues correlates with cancer aggressiveness. The intensity of the 1584 cm^−1^ Raman signal corresponding to concentration of reduced cytochrome c increases with increasing cancer aggressiveness. It indicates that cytochrome c plays a crucial role in the development and progression of cancer. The dependence of the Raman biomarker I_1584_ of the reduced cytochrome c in Fig. 3 vs cancer malignancy shows that the optimal concentration of cytochrome c in the tissue that is needed to maintain cellular homeostasis corresponds to the Raman normalized intensity of 0.006± 0.003 for the breast tissue and 0,074±0,005 for brain tissue. The concentrations of the reduced cytochrome c at this level modulate protective, signalling-response pathways, resulting in positive effects on life-history traits. The reduced cytochrome c level above the value corresponding to G0 triggers a toxic runaway process and aggressive cancer development. The plot in Fig. 3 provides an important cell-physiologic response. Normally, concentration of the reduced cytochrome c operates at low, basal level in normal cells, but it dramatically increases to very high levels in pathological cancer states.

### 2. Cytochromes in cancer human single cells

To understand this cell-physiologic response of cytochrome c due to cancer aggressiveness we used model systems of culturing lines of breast and brain cancer cells. It has become well known that the model systems of culturing cancer cells may not perfectly reproduce the biochemistry or physiology of human cancers and tumors in tissues^40^. However, they can provide information about the role of cytochrome c in single isolated cells by excluding cell-cell interactions and the effect originating from the extracellular matrix existing in the tissue.

We studied normal in vitro breast cells (MCF10A) (G0), slightly malignant MCF7 cells (G1-G2) and highly aggressive MDA-MB-231 cell (G3) and human brain cells of normal astrocytes (NHA) (G0), astrocytoma (CRL-1718) (G1-G2), glioblastoma (U-87 MG) (G4) and medulloblastoma (Daoy) (G4). We wanted to check if the main cytochrome c Raman biomarkers are also upregulated in cancers and increase with cancer aggressiveness.

To learn about cytochrome c in mitochondria or in cytoplasm by methods of conventional molecular biology one have to disrupt a cell to break it open and release the cellular structure to recover fractions that are enriched with mitochondria. Using Raman imaging we do not need to disrupt cells to learn about localization, distribution and biochemical composition of cytochromes in different organelles.

Fig. 4 A shows the Raman image of a single cell MDA-MB-231 and U-87 MG of highly aggressive breast and brain cancer and corresponding Raman spectra. Creating Raman images by K-means cluster analysis one can analyze distribution of proteins, lipids, and cytochromes in different organelles of the cell and learn about the biochemical composition from the corresponding Raman spectra. The red color represents nucleus, orange color represents lipid droplets, green-mitochondria, blue-cytoplasm, and grey-membrane.

**Figure 4.**
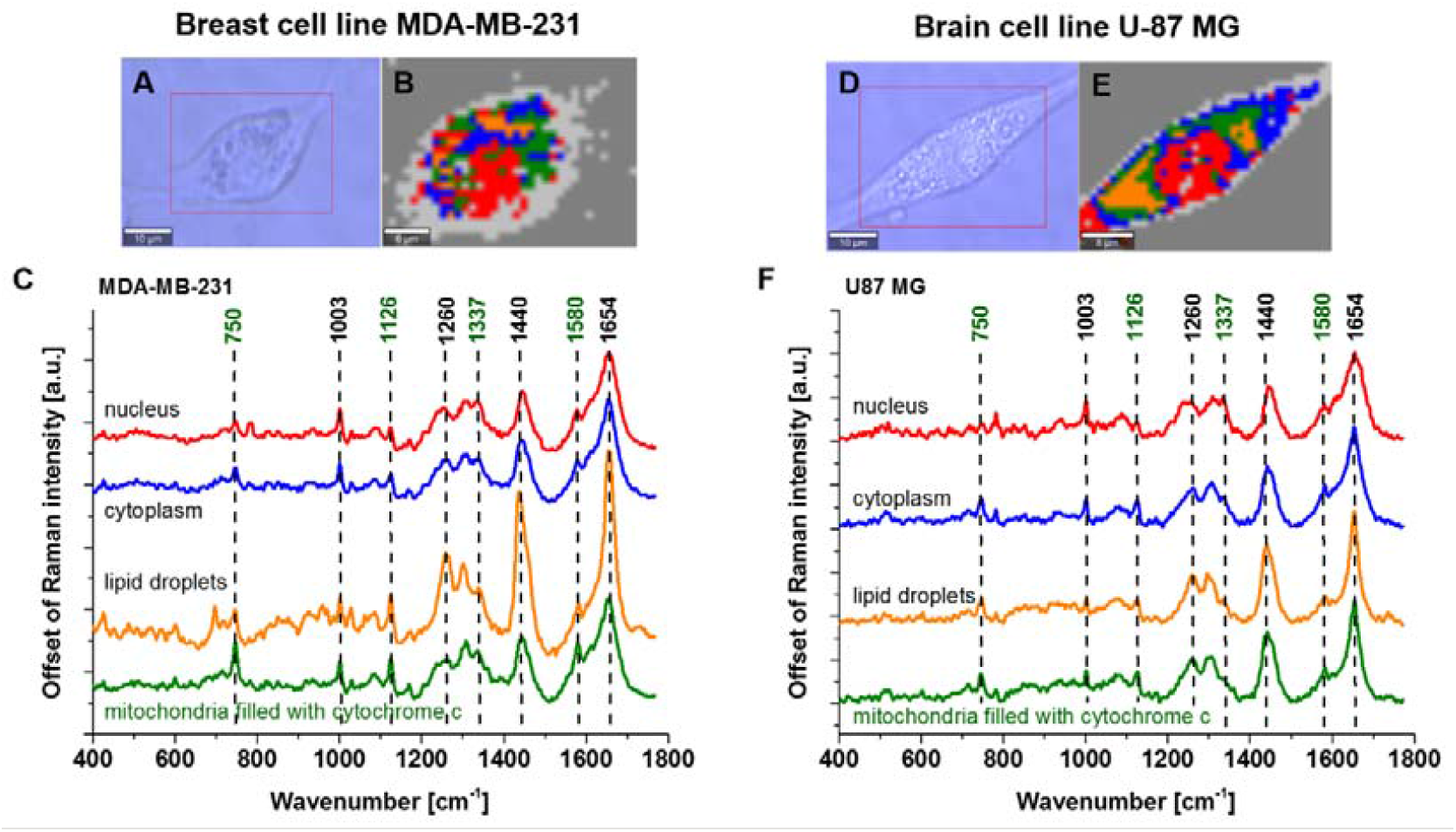
Microscopy image (A), Raman image (35×26 µm, resolution 0.5 µm, integration time 1.0 sec) (B), the average Raman spectra (number of cells n=3, number of single Raman spectra n=10920) of nuclei (red), cytoplasm (blue), lipid droplets (orange), mitochondria green) (C) at 532 nm of single cell MDA-MB-231 and Microscopy image (D), Raman image (40×30 µm, resolution 0.5 µm, integration time 1.0 sec) (E), the average Raman spectra (number of cells n=3, number of single Raman spectra n=14400) of nuclei (red), cytoplasm (blue), lipid droplets (orange), mitochondria green) (F) at 532 nm of single cell U-87 MG.

The peaks at 750 cm^−1^, ∼1126 cm^−1^ and ∼1584 cm^−1^ are associated with cytochrome c, the peak at 1337 cm^−1^ is associated with cytochrome b. Comparing the Raman intensities in different organelles of single cells one can see from Fig. 4 that the highest concentration of cytochrome c (1584 cm^−1^, 750 cm^−1^) is observed in mitochondria. We can also observe release of cytochrome c into the cytoplasm.

The significance of mitochondrial dysfunctionality has not studied by Raman method in invasive ductal carcinoma (IDC) to the best of our knowledge, but the other conventional biological method have been used to study this subject.^19^ The results on the role of cytochrome c in brain disorders of different groups are somewhat conflicted regarding the respiratory chain in glioma.^15,16^

Let us concentrate on cytochrome c concentration in mitochondria as a function of cancer aggressiveness in breast and brain cells.

Fig. 5 shows the Raman intensities of the I_1584,_ I_1126,_ I_1337,_ I_750_ cm^−1^ of vibrational peaks as a function of breast aggressiveness.

**Figure 5.**
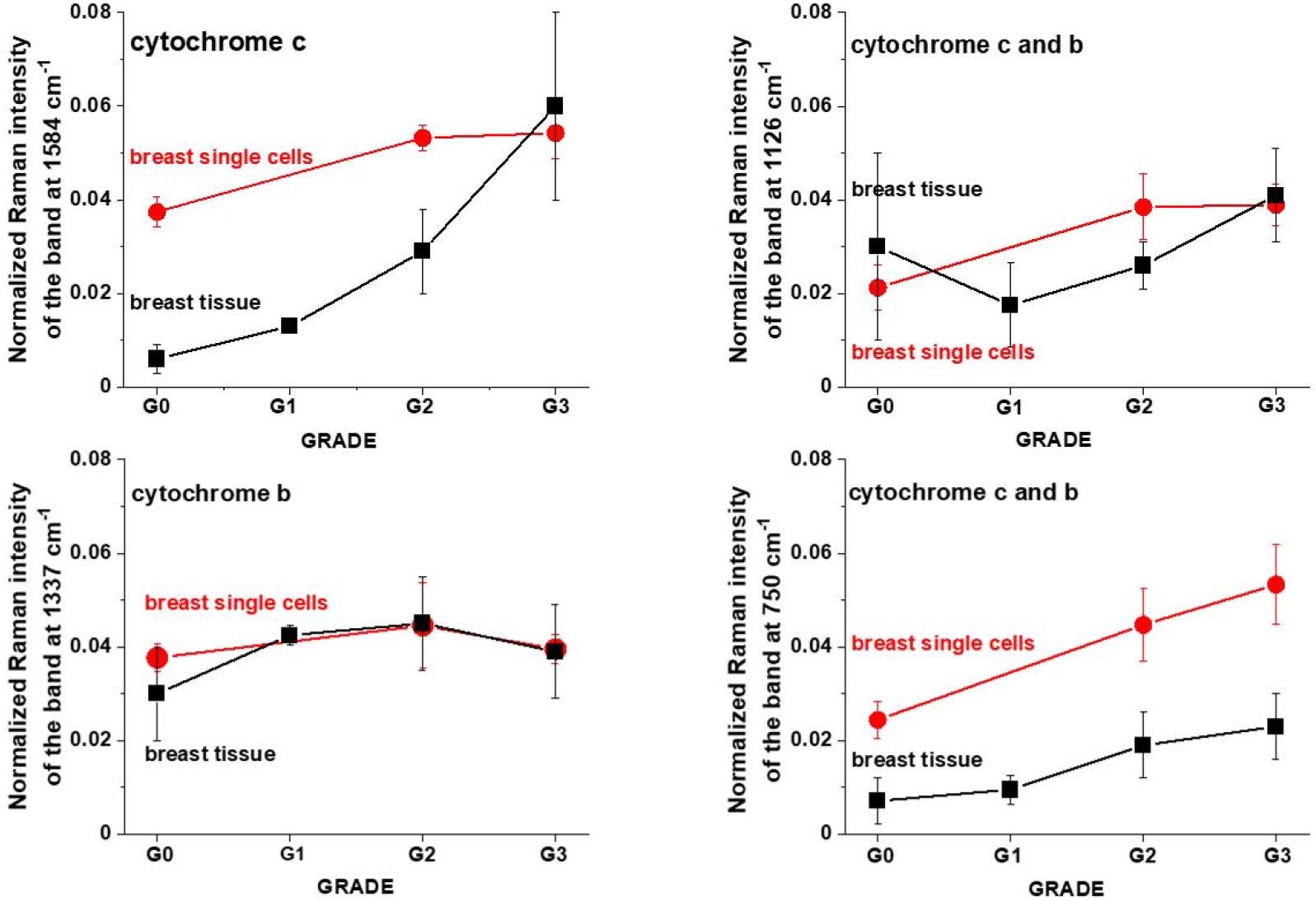
The Raman intensities of cytochrome c and cytochrome b in mitochondria of single cells in vitro culturing and in breast tissues: I_1584_, I_1126_, I_1337_, I_750_ as a function of breast cancer grade malignancy G0-G3 at excitation 532 nm.

One can see from Fig. 5 that the intensity of Raman biomarker at 1584 cm^−1^ corresponding to concentration of cytochrome c in mitochondria of a single cell increases with breast cancer aggressiveness. The intensity of Raman biomarker at 1337 cm^−1^ corresponding to concentration of cytochrome b does not change with breast cancer aggressiveness. The intensities of Raman biomarkers at 750 cm^−1^ and 1126 cm^−1^ corresponding to concentration of cytochromes c and b increase with breast cancer aggressiveness. Figure 5 demonstrate that both breast cancer tissue and breast cancer cell lines in vitro show similar trends. The higher concentration of the reduced cytochromes c in mitochondria of cancer cells (MCF7 (G2) and MDA-MB-231 (G3)) in vitro when compared with the normal cells (MCF10A (G0)) as presented in Fig. 5 indicates that the reduced form of cytochrome c is upregulated in breast cancer cells.

Fig. 6 shows the Raman intensities of the I_1584,_ I_750,_ I_1126,_ I_1337_ cm^−1^ of vibrational peaks as a function of brain aggressiveness.

**Figure 6.**
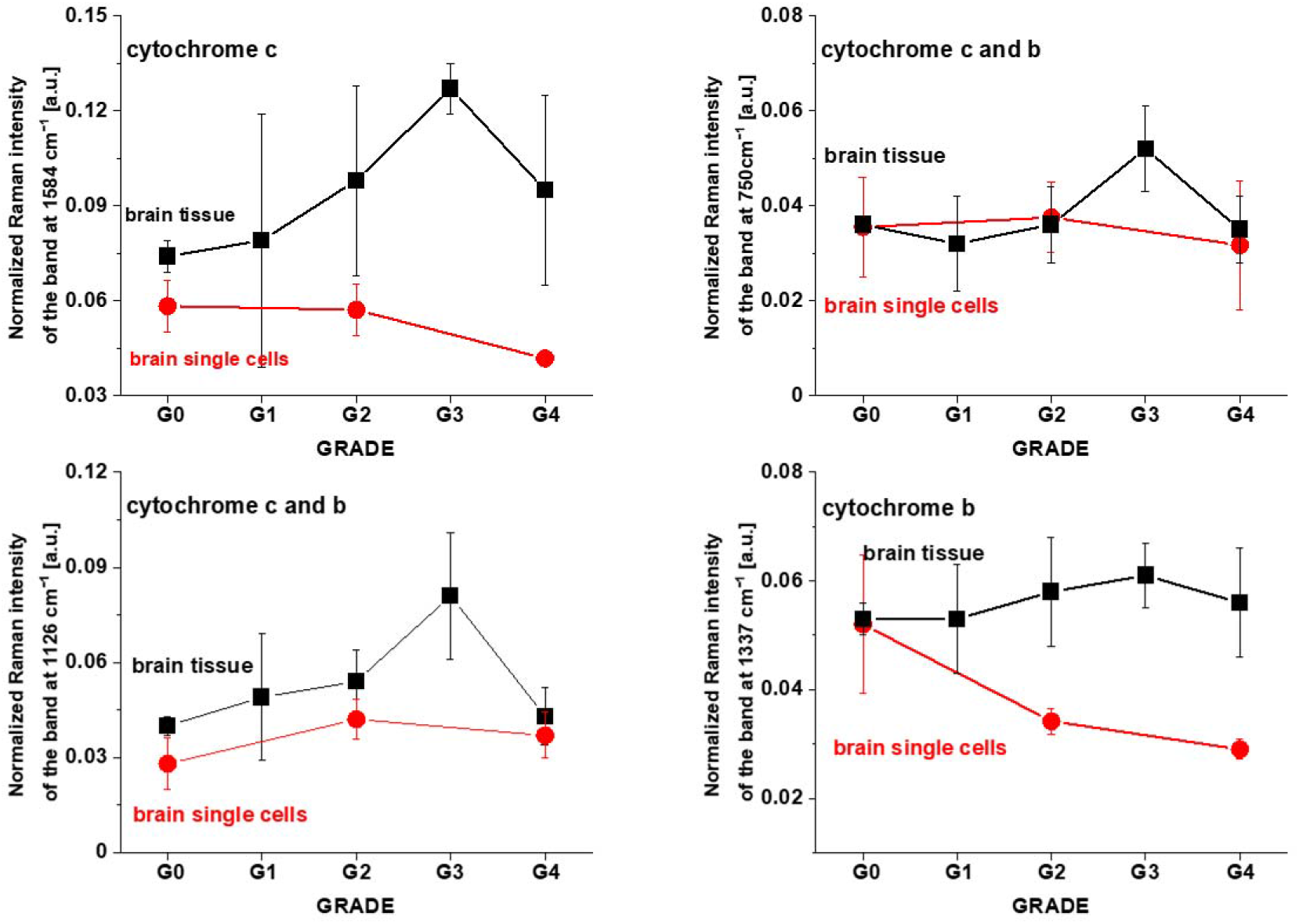
Raman intensities of cytochrome c and cytochrome b in mitochondria of single vitro cells and in the brain tissue: I_1584_, I_750,_ I_1126,_ I_1337_ as a function of brain tumor grade malignancy G0-G4 at excitation 532 nm.

One can see from Fig. 6 that the intensity of Raman biomarker at 1584 cm^−1^ corresponding to concentration of cytochrome c in mitochondria of a single cell decreases with brain tumor aggressiveness. The intensity of Raman biomarker at 1337 cm^−1^ corresponding to concentration of cytochrome b also decreases with brain tumor aggressiveness. Figure 6 demonstrate that brain tumor tissue and brain tumor single cells in vitro show opposite trends. The lower concentration of the reduced cytochromes c in mitochondria of tumor cells in vitro when compared with the normal cells as presented in Fig. 5 indicates that the reduced form of cytochrome c is downregulated in brain tumor cells.

In normal cells cytochrome c is found in the mitochondria. The release of cytochrome c into the cytoplasm induces the non-inflammatory process of apoptosis. When it is transferred to the extracellular space, it can cause inflammation. The assessment of cytochrome c in the extracellular space might be used as a biomarker for determine mitochondrial damage and cell death.

We studied concentration of cytochrome c and b using Raman markers I_750_, I_1126_, I_1584_ in cytoplasm as a function of cancer aggressiveness. Fig. 7 shows normalized Raman intensities of cytochrome c and cytochrome b in cytoplasm of single vitro cells: I_750_, I_1126_, I_1584_ as a function of breast and brain tumor malignancy at excitation 532 nm.

**Figure 7.**
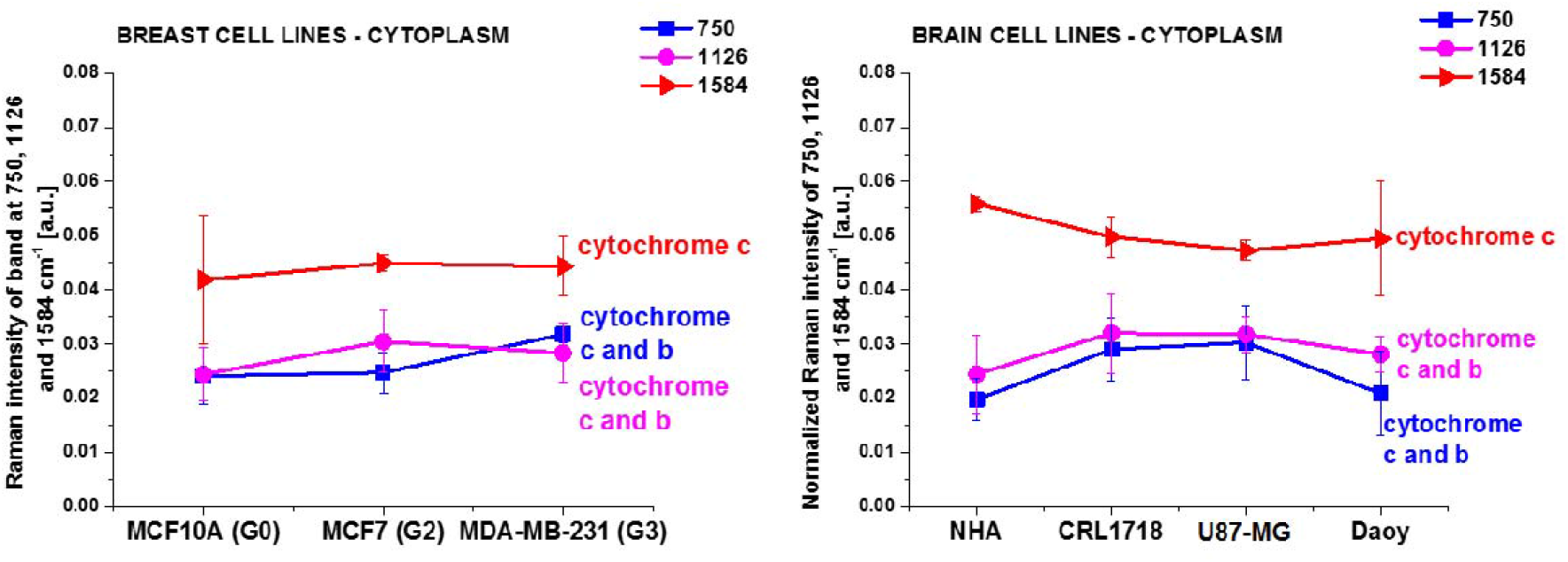
Normalized Raman intensities of cytochrome c and cytochrome b in cytoplasm of single vitro cells: I_750_, I_1126_, I_1584_ as a function of breast and brain tumor malignancy at excitation 532 nm.

One can see from Fig. 7A that in the breast single cells in vitro the concentration of cytochrome c found in the cytoplasm slightly increases with cancer aggressiveness. The release of cytochrome c into the cytoplasm induces the non-inflammatory process of apoptosis. The effect is much stronger in breast cancer tissue as one can see in Fig. 6 where we observe the mixed effect of the release of cytochrome c into the cytoplasm and extracellular space where it can cause inflammation. From Fig. 7B one can see that in the brain single cells in vitro the concentration of cytochrome c found in the cytoplasm is less clearly visible vs. cancer aggressiveness than in breast cancer.

Recent studies have confirmed the release of cytochrome c into the extracellular space and ultimately into the blood in various conditions characterizes cell death. Obviously, the greater the damage of cells or tissues, the higher the serum cytochrome c level. Thus, cytochrome c may be a useful clinical biomarker for diagnosing and assessing pathological entities. It suggests that cytochrome c level in serum after chemotherapy may be a good prognostic factor, indicating an increased apoptosis of cancer cell induced by chemotherapy.^41-44^

### 3. Discrepancies between tissues and in vitro cells vs cancer aggressiveness

In the view of the results presented so far shows that breast cancer tissue and breast cancer cell lines in vitro show similar trends. The concentration of cytochrome c increases with increasing breast cancer aggressiveness. In contrast, the results demonstrate that brain tumor tissue and brain tumor single cells in vitro show opposite trends. The concentration of cytochrome c increases with increasing brain tumor aggressiveness in the tissue, while the concentration of cytochrome c decreases with increasing breast cancer aggressiveness in single cells in vitro. Here, we will discuss the possible reason of these discrepancies between tissues and in vitro cells vs cancer aggressiveness to explain why this relationship is reversed. First, in single cells we analyze concentration of cytochromes in separate organelles like mitochondria in Figures 5 and 6. In the tissue we measure the global concentration of cytochrome, c not only inside the cell, but also outside in the extracellular matrix due to the release of cytochrome from cells. Second, interactions between epithelial cells and extracellular matrix in the tissue results in induction of various mechanisms like caspase-dependent cytochrome c release that may occur by distinct mechanisms than in single cells in vitro.

To clarify this point let us focus on the role of cytochrome c in mechanisms that regulate mitochondrial dysfunction in cancers. Our results for single cells presented above provide a first hint on the role of cytochrome c in mechanisms that regulate cancer progression by using redox-sensitive mitochondrial cytochrome Raman bands for label-free detection of mitochondrial dysfunction.

To understand the role of cytochrome c let’s summarize the most important events with its involvement in oxidative phosphorylation (OXPHOS). Briefly, as shown in Scheme 1A, a series of coordinated enzymatic reactions are involved in the flow of carbons from glucose to fatty acids. First, glucose derived from dietary carbohydrates undergoes glycolysis, pyruvate generated from glycolysis is changed into the compound known as acetyl CoA. Many intermediate compounds are formed in the TCA cycle which are used in synthesis of other biomolecules, such as amino acids, nucleotides, chlorophyll, cytochromes and fats. The third part of cellular respiration in mitochondria is oxidative phosphorylation which is essential to the maintenance of life. OXPHOS process oxidises glucose derivatives, fatty acids and amino acids to carbon dioxide (CO_2_) through a series of enzyme controlled steps. In this part OXPHOS starts with the respiratory electron transport chain (Scheme 1B) transferring electrons originating from NADH and succinate through four protein complexes (complex I ∼ IV) with the help of ubiquinone and cytochrome c. During the electron transfer process, a proton gradient is generated across the inner mitochondrial membrane, which in turn drives ATP synthase to synthesize ATP.^45^

**Scheme 1.**
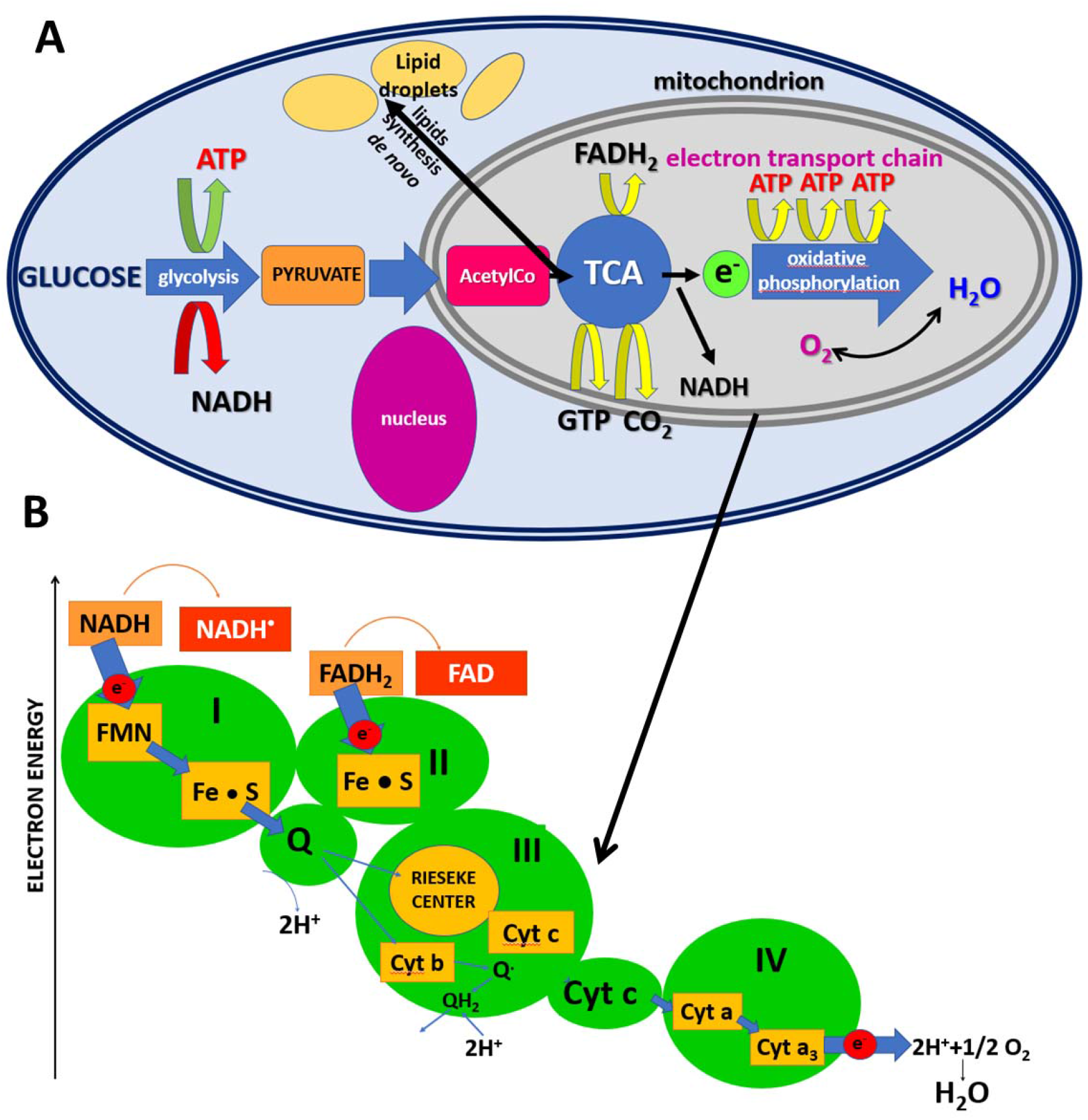
Glycolysis takes place in the cytoplasm. Within the mitochondrion, the citric acid cycle (TCA) occurs in the mitochondrial matrix, and oxidative metabolism and the electron transport chain occurs at the internal mitochondrial membranes.

Scheme 1B shows the electron transport chain with involvement of cytochrome c. Briefly, the electrons are transported in a series of events via electron transporters embedded in the inner mitochondrial membrane that shuttles electrons from NADH and FADH_2_ to molecular oxygen. As a result of these processes, protons are pumped from the mitochondrial matrix to the intermembrane space, and oxygen is reduced to form water. The cytochrome c is localized in the inner membrane of mitochondrium and transfers electrons between complex III and complex IV of the respiratory chain (Scheme 1) generating the proton gradient and triggering ATP production. The heme chromophore of cytochrome c accepts electrons from the bc_1_ complex (complex III) and transfers electrons to the complex IV. Under normal physiological conditions, when the electron transport chain and chemiosmosis make up oxidative phosphorylation, there is a balance between the oxidized and reduced forms of cytochrome c, which depends on the efficiency of the electron transport from complex I to complex IV.

Our results presented so far demonstrate that the mechanisms controlling the electron transport chain are spectacularly deregulated in cancers and the level of dysfunction measured as concentration of reduced cytochrome c increases with tumor aggressiveness. At normal physiological conditions the oxidized form of the cytochrome c heme group can accept an electron from the heme group of the cytochrome c1 subunit of cytochrome reductase (complex III). Cytochrome c then transfers this electron to the cytochrome oxidase complex (complex IV).

Our results in Fig. 6 demonstrate that concentration of reduced cytochrome c in mitochondria of brain single cells monitored by the Raman signal at 1584 cm^−1^ decreases with increasing malignancy level. It indicates that complex III shows reduced activity in transferring electron to cytochrome c with increasing malignancy level. Additionally, concentration of cytochrome b also decreases with tumor malignacy (Fig. 6 C). The results from Fig. 6 C suggests that cancer cells are deficient in subunit cytochrome b in the complex III, which are unable to maintain respiratory function. Thus, the results from Fig. 6 demonstrate that electron transport, organized in terms of electronegativity, is inhibited between complex III and complex IV (Scheme 1).

The results for brain support earlier suggestions that the Q_o_ site of the mitochondrial complex III is required for the transduction of hypoxic signalling via reactive oxygen species production.^46^ Cancer cells deficient in subunit cytochrome b in the complex III, which are unable to maintain respiratory function, increase ROS levels and stabilize the HIF-1α protein during hypoxia.^46^ CYC1 is a phosphoprotein and subunit of ubiquinol cytochrome c reductase that binds heme groups.^46^

The mechanism of oxidative phosphorylation with involvement of cytochrome c in breast cancer seems to be a bit different than in brain tumors. Indeed, the large pool of reduced cytochrome c that increase with cancer aggressiveness (Fig. 5 A, B) suggests that the origin of mitochondrial dysfunction comes from complex IV, the last enzyme in the respiratory electron transport chain of cells. Thus, in contrast to brain tumors the results for breast cancer would rather suggest dysfunction of the complex IV.

The complex IV contains two hemes, cytochrome a and cytochrome a_3_, and two copper centers, the Cu_A_ and Cu_B_ centers and several subunits belonging to COX family. Complex IV receives an electron from each of four cytochrome c molecules, and transfers them to one dioxygen molecule, converting the molecular oxygen to two molecules of water. In this process it binds four protons from the inner aqueous phase to make two water molecules, and translocates another four protons across the membrane, increasing the transmembrane difference of proton electrochemical potential which triggers the ATP synthase to provide energy. In addition to providing energy, cytochrome c has other essential role within cells: it is one of regulators of biosynthesis in lipid synthesis de novo.

### 4. Lipid synthesis de novo

It is known that certain cytochromes such as P450 enzymes (CYP) are critical in metabolizing polyunsaturated fatty acids (PUFAs) to biologically active, intercellular cell signaling molecules (eicosanoids) and/or metabolize biologically active metabolites of the PUFA to less active or inactive products. These CYPs possess cytochrome P450 omega hydroxylase and/or epoxygenase enzyme activity.^47^

It is possible that cyclooxygenase (COX) overexpression observed in cancers^48,49^ is related to disruption in the process of electron transfer from cytochrome c. Detailed analysis will be necessary to find correlation between conformations and other alterations in COX subunits and electron transfer from cytochrome c. Since COX inhibitors belong to the most commonly taken drugs^50,51^, further research should focus on understanding the mechanisms of correlation. The origin of mitochondrial dysfunction of complex IV in cancers is still unknown, but our previous results demonstrated that there is a link between lipid reprogramming and COX family^34^ in breast cancerogenesis. These observations led us to hypothesize a role for the cytochrome family in mechanisms of lipid reprogramming that regulates cancer progression.

To better understand the link between lipid metabolism and mitochondrial function of cytochrome c let’s look once again at the main pathways described in the Scheme 1A. Pyruvate generated from glycolysis is changed into the compound known as acetyl CoA. The acetyl CoA enters tricarboxylic acid (TCA) cycle resulting in a series of reactions. The first reaction of the cycle is the condensation of acetyl-CoA with oxaloacetate to form citrate, catalyzed by citrate synthase. One turn of the TCA cycle is required to produce four carbon dioxide molecules, six NADH molecules, and 2 FADH_2_ molecules. TCA cycle occurs in the mitochondria of the cell. Citrate from the TCA cycle is transported to cytosol and then releases acetyl-CoA by ATP-citrate lyase (ACLY). The resulting acetyl-CoA is converted to malonyl-CoA by acetyl-CoA carboxylases. Then, fatty acid synthase (FASN), the key rate-limiting enzyme in de novo lipogenesis (DNL), converts malonyl-CoA into palmitate, which is the first fatty acid product in DNL. Finally, palmitate undergoes the elongation and desaturation reactions to generate the complex fatty acids, including stearic acid, palmitoleic acid, and oleic acid, which we can observe by Raman imaging as lipid droplets (LD). We showed the lipid droplets are clearly visible in Raman images and we analysed the chemical composition of LD in cancers.^6,52^

Fig. 8 shows the normalized Raman intensities 1444 cm^−1^ corresponding to vibrations of lipids in human normal and cancer tissues and in lipid droplets in single cells in vitro as a function of cancer grade malignancy at excitation 532 nm.

**Figure 8.**
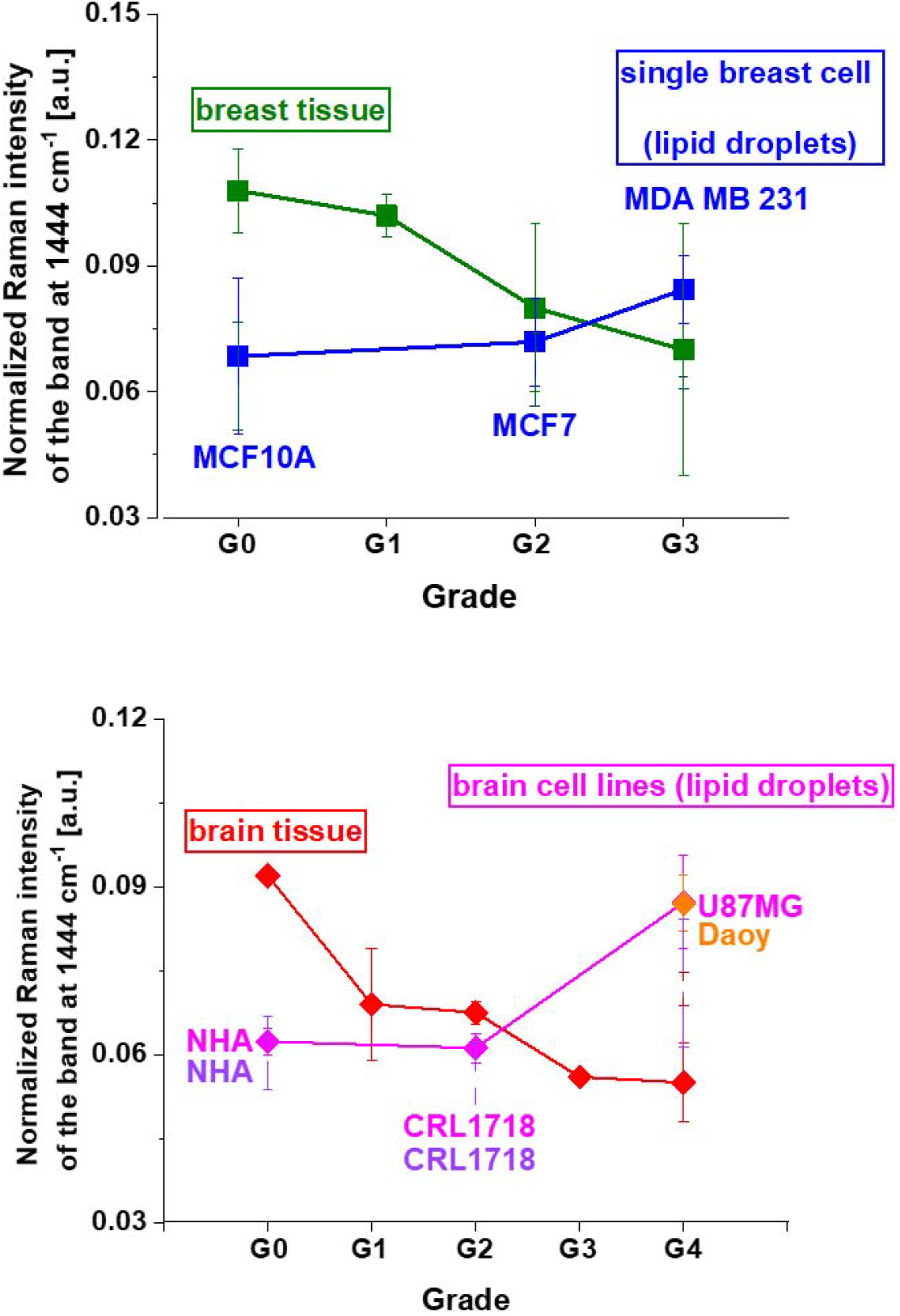
The normalized Raman intensities of lipids in human normal and cancer tissues and cells in vitro at 1444 cm^−1^ as a function of breast (A) and brain (B) cancer grade malignancy at excitation 532 nm.

One can see that the intensity of the band at 1444 cm^−1^ increases with cancer aggressiveness in lipid droplets both in breast and brain single cells in contrast to human cancer tissues. Again, like for Raman biomarkers of cytochrome presented in Figures 5 and 6, the relationship between the concentration of lipids vs aggressiveness is reversed.

To explain this finding we have to remind that lipids can be provided by diet or by de novo synthesis. While glioma or epithelial breast cells clearly rely upon fatty acids for energy production, it is not clear whether they acquire fatty acids from the bloodstream or build these carbon chains themselves in de novo lipogenesis. The answer can be provided from comparison between single cells and cancer tissue vs cancer aggressiveness. Fig. 8 shows that in breast and brain tissues the normalized Raman intensity of fatty acids at 1444 cm^−1^decreases, not increases, with increasing cancer grading in contrast to single cells. It indicates that in tissue contribution from the bloodstream dominates over de novo fatty acids production. It explains discrepancies between lipids level in tissues and in vitro cells vs cancer aggressiveness presented in Fig.8.

Detailed inspection into Fig.8 demonstrates that for in vitro cell lines the enhanced lipogenesis (monitored by the Raman peak at 1444 cm^−1^) increases with breast and brain cancer aggressiveness. It is worth emphasizing that enhanced lipogenesis de novo is positively correlated with the concentration of reduced cytochrome c in single breast cells (750 cm^−1^, 1126 cm^−1^ and 1584 cm^−1^ in Fig. 5). In contrast, the enhanced lipogenesis is inversely related to the concentration of reduced cytochrome c in single brain cells (750 cm^−1^, 1126 cm^−1^ and 1584 cm^−1^ in Fig. 6). It indicates that for single cells in vitro, when the extracellular matrix and cell-cell interactions are eliminated, the lipid de novo biosynthesis increases whereas oxidative phosphorylation controlled by cytochrome c decreases with increasing cancer aggressiveness. The situation in the tissues, where the extracellular matrix and cell-cell interactions are not eliminated, the relationship is reversed (Figs.6-8).

Recently, we found increased amount of cytoplasmic lipid droplets in the human cancer cells, which must be closely related to increased activity of lipid synthesis in cancerous tissues.^52^ We showed that de novo lipogenesis is significantly enhanced in cancer cells.^9^ In de novo mechanism inside cytosol of a cell fatty acids are shuttled into lipid droplets upon hypoxia in tumors in order to support cell growth and survival upon re-oxygenation.^53^ Increasingly it is appreciated that fatty acids can act as critical bio-energetic substrates within many cancers including the glioma cell.^40^

Two types of lipid droplets in normal astrocytes and cancer cells of glioblastoma with distinct chemical compositions, biological functions and vibrational properties have been found.^52^ Two types of lipid droplets are related to different functions - energy storage and signalling. Their expression and biochemical composition depend on cancer aggressiveness. The cancer cells are dominantly filled with TAGs and are involved in energy storage. The normal cells are mainly filled with retinyl esters and retinol binding proteins and are involved in signalling, especially JAK2/STAT6 pathway signalling. The TAG are dominated by polyunsaturated fatty acids (PUFAs) identified as arachidonic acid esters (AA).^54^ Cyclooxygenases COX catalyzes the conversion of the free essential fatty acids to prostanoids. Prostanoids represent a subclass of eicosanoids consisting of: the prostaglandins (mediators of inflammatory and anaphylactic reactions), the thromboxanes (mediators of vasoconstriction) and the prostacyclins (active in the resolution phase of inflammation).^34^

The deficiency of complex IV containing COX units and related to electron transfer along complex III-cytochrome c-complex IV may control and enhance inflammatory processes that lead to cancer development.

Our results allow to look from a new perspective on the triangle between altered bioenergetics, enhanced biosynthesis and redox balance in cancer development.

To check the shift in glucose metabolism from oxidative phosphorylation to lactate production for energy generation (the Warburg Effect), a well-known metabolic hallmark of tumor cells, we used the Raman peak at 823 cm^−1^ presented in Fig. 9 to detect the presence of the lactic acid.

**Figure 9.**
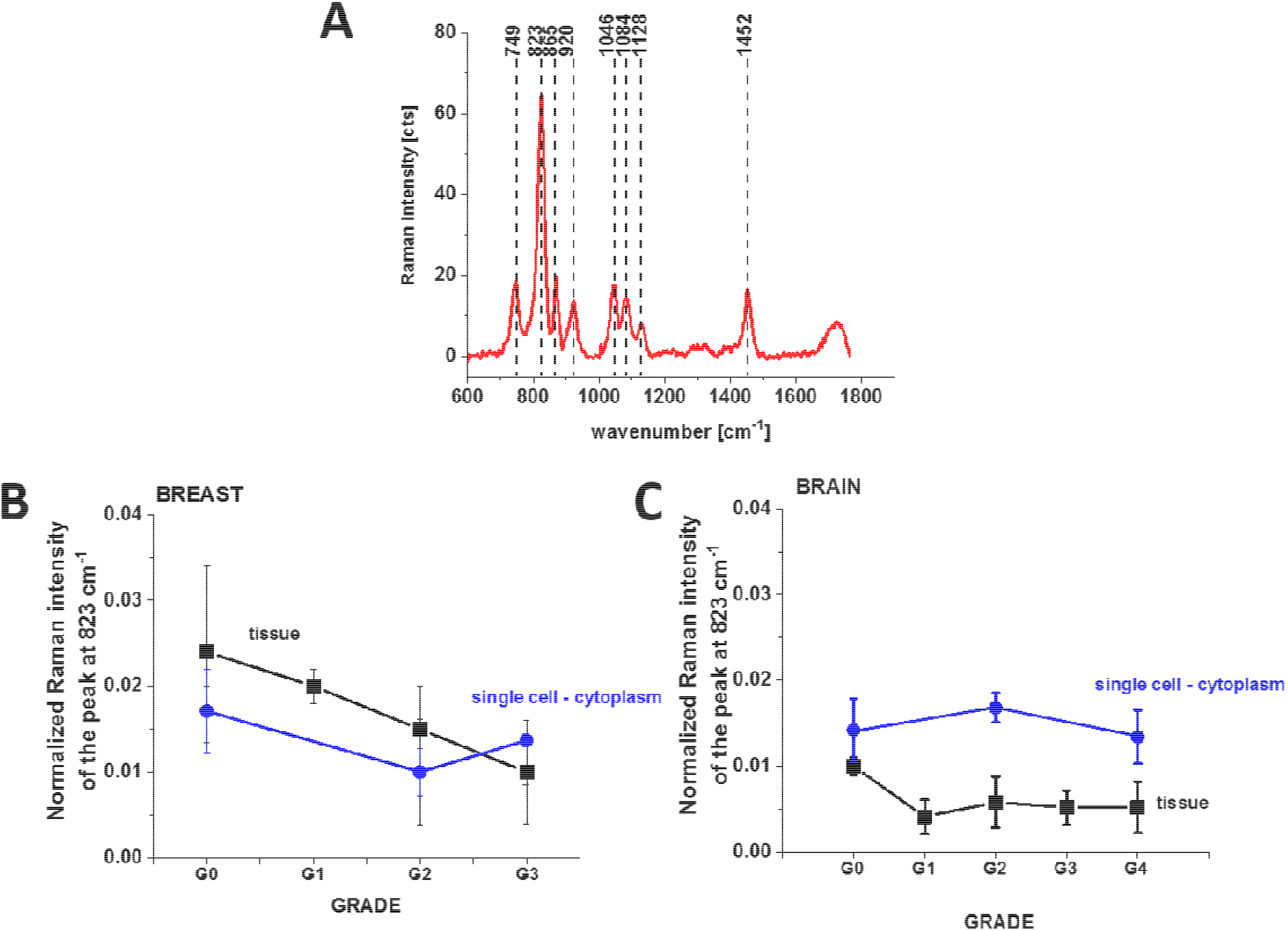
Raman spectrum of lactic acid (A), Raman intensities of peak 823 as a function of human tissue breast cancer malignancy (G1-G3) (B) and of human tumor brain malignancy (G1-G4) (C) excitation 532 nm.

One can see from Fig.9 that the Raman intensity of the band 823 cm^−1^ corresponding to the concentration of lactic acid in breast and brain single cells in cytoplasm and in tissues decreases, not increases, vs cancer aggressiveness, when compared with the normal brain and breast tissues. It indicates that the efficiency of the switch in glucose metabolism from oxidative phosphorylation to lactate production decreases with cancer aggressiveness. These results combined with the results presented in Figures 5 show that metabolic adaptation in tumors extends beyond the Warburg effect. Indeed, the results from Figures 5 show that concentration of one of the most important molecule in oxidative phosphorylation - cytochrome c - in mitochondria increases with breast cancer aggressiveness.

The results suggest that the metabolic adaptation in tumors follow the same pattern of behavior as in normal cells by inducing mechanism of higher cytochrome c concentration to maintain oxidative phosphorylation. The path of oxidative phosphorylation is needed to maintain enhanced biosynthesis including ATP and de novo fatty acids production. We showed that de novo fatty acids production detected by the Raman intensity at 1444 cm^−1^ increases with cancer aggressiveness in contrast to production of lactic acid detected by the Raman intensities at 823 cm^−1^ that decreases with cancer aggressiveness for single cancer cells in vitro. Based on the Raman intensities of the vibrations corresponding to cytochrome c, fatty acids, and lactic acid we discovered that in breast cancer cells that total ATP turnover was 75% oxidative and 25% glycolytic. Presently, an increasing number of reports have supported our results about metabolic regulation in cancers^40,55,56^ showing that metabolic adaptation in tumors are highly oxidative. Recently, it was discovered that in MCF-7 breast cancer cells that total ATP turnover was 80% oxidative and 20% glycolytic.^57^ This hypothesis was also tested in primary-cultured human glioblastoma cells and it was found that cells were highly oxidative and largely unaffected by treatment with glucose or inhibitors of glycolysis.^5^ Thus, it appears that oxidative phosphorylation can not only co-exist with aerobic glycolysis and lactate release, but it dominates metabolic adaptation in tumors.

## Conclusions

The results suggest that Raman spectroscopy, in the future, could be an alternative technique for monitoring the relations between altered bioenergetics, enhanced biosynthesis and redox balance in cancer development. Our results suggest that the shift in glucose metabolism from oxidative phosphorylation to lactate production for energy generation (the Warburg Effect), a well-known metabolic hallmark of tumor cells, is not dominant mechanism of cancer development. Our results show that the cancer cells follow the same pattern of behavior as normal cells by inducing mechanisms of higher cytochrome c concentration to maintain oxidative phosphorylation in the electron transport chain required to fuel bioenergetics via ATP and enhance de novo biosynthesis of lipids. The Warburg effect by converting glucose to lactate is only additional mechanism, which is far less efficient in ATP production than oxidative phosphorylation. The efficiency of Warburg mechanism decreases with increasing tumor aggressiveness. Based on the Raman intensities of the vibrations corresponding to cytochrome c, fatty acids, and lactic acid we discovered that in breast cancer cells that total ATP turnover was 75% oxidative and 25% glycolytic.

We showed that Raman imaging provides additional insight into the biology of gliomas and breast ductal invasive cancer, which can be used for non-invasive grading, differential diagnosis, delineation of tumor extent, planning of surgery, and radiotherapy and post-treatment monitoring. We used Raman spectroscopy to monitor changes in the redox state of the mitochondrial cytochromes in ex vivo human brain and breast tissues surgically resected specimens of human and, in vitro human brain cells of normal astrocytes (NHA), astrocytoma (CRL-1718), glioblastoma (U87-MG) and medulloblastoma (Daoy), and human breast cells of normal cells (MCF 10A), slightly malignant cells (MCF7) and highly aggressive cells (MDA-MB-231) at 532 nm.

Our results show that human breast and brain cancers demonstrate a redox imbalance compared to normal tissues. The reduced cytochrome c is upregulated in cancers. The results of the paper shed light on a largely non-investigated triangle between cytochromes, lipid metabolism and mitochondrial function in electron transfer chain. The results presented in the paper providing insight into the crosstalk between organelles increases our understanding of mitochondria-driven cancer. In the paper we explored a hypothesis involving the possible role of redox state of cytochrome c in cancer. We found biochemical modifications in cellular mitochondria, lipid droplets and cytoplasm observed in cancer progression which are caused by redox imbalance.

The biochemical results obtained by Raman imaging showed that human single cells in vitro demonstrate a redox imbalance by upregulation of cytochrome c in breast ductal cancer and downregulation of cytochrome c in brain tumors. Both breast and brain tumors demonstrate enhanced lipogenesis de novo compared to normal cells.

The paper demonstrates the critical role of extracellular matrix in mechanisms of oxidative phosphorylation. We showed that concentration of reduced cytochrome c (monitored at 1584 cm^-1^) is lower in cancer single cells when comparted with the normal cells at in vitro conditions when the effect of microenvironment is eliminated. In contrast, the redox balance shows reverted trend in the breast cancer and brain tumor tissues when there are interactions with the environment. The concentration of reduced cytochrome c (monitored at 1584 cm^-1^) is significantly higher in cancer tissue when comparted with the normal tissue.

Our results suggest that the mechanisms controlling the electron transport chain are spectacularly deregulated in cancers. The electron transport, organized in terms of electronegativity, is inhibited between complex III and cytochrome c for isolated breast cells in vitro and between cytochrome c and complex IV in brain cells. This study demonstrates the ability of confocal Raman microscopy to detect apoptosis mediated by cytochrome c release from mitochondria.

The results presented in the paper suggest that the redox-sensitive peak observed at 1584 cm^−1^ with excitation at 532 nm is specifically linked to cytochrome c and can be considered to be a “redox state marker” of the ferric low spin heme in cyt c, assigned to the v_19_ mode, vibrations of methine bridges (C_α_C_µ_, C_α_C_µ_H bonds) and the C_α_C_β_bond.

Our results show that cytochrome c concentration correlates with cancer aggressiveness. The higher concentration of cytochrome c demonstrates high-turnover and more aggressive tumors. Obviously, the greater the damage of cells or tissues, the higher the serum cytochrome c level. Thus, cytochrome c may be a useful clinical biomarker for diagnosing and assessing pathological entities. The results presented in the paper may provide a new opportunity in cancer prevention and treatment that involves cytochrome family. However, further studies are required for supporting this role for cytochrome c and the responsible pattern recognition receptors remain to be discovered.

## Materials and Methods

### Reference chemicals

Cytochrome c (no. C2506), DL-lactic acid (no. 69785) were purchased from Sigma Aldrich.

### Ethics statement

All the conducted studies were approved by the local Bioethical Committee at the Polish Mother’s Memorial Hospital Research Institute in Lodz (53/216) and by the institutional Bioethical Committee at the Medical University of Lodz, Poland (RNN/323/17/KE/17/10/2017). Written consents from patients or from legal guardians of patients were obtained. All the experiments were carried out in accordance with Good Clinical Practice and with the ethical principles of the Declaration of Helsinki. Spectroscopic analysis did not affect the scope of surgery and course and type of undertaken hospital treatment.

### Patients

In the presented studies the total number of patients diagnosed with brain tumors was 44. Among them 11 were diagnosed with medulloblastoma, 1 with embryonic tumor PNS, 3 with ependynoma anaplastic, 4 with ependymoma, 2 with astrocytoma fibrous, 1 with astrocytoma, 1 with ganglioma, 8 with astrocytoma pilocytic, 1 with subependymoma, 2 with hemangioblastoma, 4 with craniopharyngioma, 1 with dysembryoplastic neuroepithelial tumor, 1 with papillary glioneuronal tumor, 1 with testicular cancer metastasis, 1 with gliosarcoma, 1 with anaplastic oligodendroglioma, and 1 sample it was tumor metastasis. All patients were treated at the Polish Mother’s Memorial Hospital Research Institute in Lodz. For breast cancers the number of patients was 39, all patients were diagnosed with infiltrating ductal carcinoma and treated at the M. Copernicus Voivodeship Multi-Specialist Center for Oncology and Traumatology in Lodz.

### Tissues samples collection and preparation for Raman spectroscopy

Tissue samples were collected during routine surgery. The non-fixed samples were used to prepare 16 micrometers sections placed on CaF_2_ substrate for Raman analysis. In parallel typical histopathological analysis by professional pathologists from the Polish Mother’s Memorial Hospital Research Institute in Lodz for brain tissues samples or from Medical University of Lodz, Department of Pathology, Chair of Oncology for breast tissues samples was performed. The types and grades of tumors according to the criteria of the Current WHO Classification were diagnosed.

### Cell culture and preparation for Raman spectroscopy

The studies were performed on a normal human astrocytes (Clonetics NHA), human astrocytoma CCF-STTG1 (ATTC CRL-1718) and human glioblastoma cell line U87-MG (ATCC HTB-14) purchased from Lonza (Lonza Walkersville. Inc.) and American Type Culture Collection (ATCC), respectively. The NHA cells were maintained in Astrocyte Medium Bulletkit Clonetics (AGM BulletKit, Lonza CC-3186) and ReagentPack (Lonza CC-5034) without antibiotics in a humidified incubator at 37°C and 5% CO_2_ atmosphere. The U87MG cells were maintained in Eagle’s Minimal Essential Medium Eagle with L-glutamine (ATCC 30-2003) supplemented with 10% fetal bovine serum (ATCC 30-2020) without antibiotics in a humidified incubator at 37°C and 5% CO_2_ atmosphere. The CRL-1718 cells were maintained in RPMI1640 Medium (ATCC 30-2001) supplemented with 10% fetal bovine serum (ATCC 30-2020) without antibiotics in a humidified incubator at 37°C and 5% CO_2_ atmosphere. A human desmoplastic cerebellar medulloblastoma cell line (ATCC HTB-186, Daoy) was grown in Eagle’s Minimum Essential Medium (EMEM, ATCC 30-2003) supplemented with the fetal bovine serum to a final concentration of 10% (Gibco, Life Technologies, 16000-044). Cells were maintained without antibiotics at 37°C in a humidified atmosphere containing 5% CO_2_. A human breast MCF10A cell line (CRL10317, ATCC) was grown with completed growth medium: MEGM Kit (Lonza CC3150) without gentamycin-amphotericin B mix (GA1000) and with 100 ng/ml cholera toxin; a slightly malignant human breast MCF7 cell line (HTB22, ATCC) in Eagle’s Minimum Essential Medium (ATCC 30-2003) with 10% fetal bovine serum (ATCC 30-2020) and highly aggressive human breast MDA-MB-231 cell line (HTB26, ATCC) in Leibovitz’s L15 Medium (ATCC 30-2008) with 10% fetal bovine serum (ATCC 30-2020). All human breast cell lines were maintained at 37°C in a humidified atmosphere containing 5% CO_2_. Cells were seeded on CaF_2_ window in 35mm Petri dish at a density of 5×10^4^ cells per Petri dish the day before examination. Before Raman examination, cells were fixed with 4% formalin solution (neutrally buffered) and kept in phosphate buffered saline (PBS, Gibco no. 10010023) during the experiment.

### Raman human tissues spectroscopic measurements *ex-vivo*

WITec (Ulm, Germany) alpha 300 RSA+ confocal microscope was used to record Raman spectra and imaging. The configuration of experimental set up was as follows: the diameter of fiber: 50 μm, a monochromator Acton-SP-2300i and a CCD camera Andor Newton DU970-UVB-353, the excitation laser line 532 nm.. Excitation line was focused on the sample through a 40x dry objective (Nikon, objective type CFI Plan Fluor C ELWD DIC-M, numerical aperture (NA) of 0.60 and a 3.6–2.8 mm working distance). The average laser excitation power was 10 mW, with an integration time of 0.5 sec. An edge filters were used to remove the Rayleigh scattered light. A piezoelectric table was used to record Raman images. No pre-treatment of the samples was necessary before Raman measurements. The cosmic rays were removed from each Raman spectrum (model: filter size: 2, dynamic factor: 10) and the smoothing procedure: Savitzky–Golay method was also implemented (model: order: 4, derivative: 0). Data acquisition and processing were performed using WITec Project Plus software.

### Statistical analysis

All results regarding the analysis of the intensity of the Raman spectra as a function of breast cancer or brain tumor grades are presented as the mean±SD, where p<0.05; SD - standard deviation, p – probability value. Raman bands intensity were taken from normalized by vector norm spectra. The Raman spectra were obtained from 39 patients (breast) and 44 (brain). For each patient thousand spectra from different sites of the sample were obtained from cluster analysis. For breast we used typically 8000 Raman spectra for averaging, and 6500 Raman spectra for brain tissue.

### Cluster analysis

Spectroscopic data were analysed using Cluster Analysis method. Briefly Cluster Analysis is a form of exploratory data analysis in which observations are divided into different groups that have some common characteristics – vibrational features in our case. Cluster Analysis constructs groups (or classes or clusters) based on the principle that: within a group the observations must be as similar as possible, while observations belonging to different groups must be as different.

The partition of n observations (x) into k (k≤n) clusters S should be done to minimize the variance (Var) according to the formula:

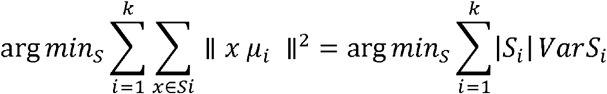

where *µ*_*i*_is the mean of points *s*_*i*_

## Acknowledgement

This work was supported by the National Science Centre of Poland (Narodowe Centrum Nauki, UMO-2019/33/B/ST4/01961). Daoy cell line was funding from the grant no 2018/02/X/NZ3/00590 (Miniatura2, ID 407029, Narodowe Centrum Nauki).

## Author Contributions

Conceptualization: HA; Funding acquisition: HA, JS; Investigation: BB-P, JS, MK; Methodology: HA, BB-P, JS, MK; Writing – original draft: HA; Writing – review & editing: HA, BB-P, JS, MK. All authors reviewed and provide feedback on the manuscripts.

